# Impact of repetition on physical and imagined movements

**DOI:** 10.64898/2026.09.22.753405

**Authors:** Elise E Van Caenegem, Marcos Moreno-Verdú, Charlène Truong, Baptiste M Waltzing, Robert M Hardwick

## Abstract

Motor imagery and overt action are proposed to share common planning mechanisms while relying on distinct processes during movement performance. To investigate how repetition affects physical and imagined movements, participants physically or mentally drew paths while avoiding barriers. The barrier orientations were either repeated or different across consecutive trials, and the modality (execution or imagery) of the trials either changed or remained the same for two consecutive trials. Reaction time and movement time were analysed as measures of movement planning and task performance, respectively.

Imagined movements produced longer reaction times and movement times than physical movements. Repetition of orientation reduced reaction times in both physical and imagined trials, indicating that trajectory repetition facilitated movement preparation. In contrast, previous trial type influenced reaction times only for physical movements, which were only initiated faster when preceded by another physical trial. Different effects emerged for movement time. Physical movements were faster when the same trajectory was repeated no matter the previous trial type, whereas imagined movements were unaffected by orientation and instead showed shorter durations when preceded by another imagined trial.

The findings indicate that physical and imagined movements rely on more similar trajectory representations during the planning stage, but diverge during task performance. While physical movements appear to benefit from the persistence of previously generated movement trajectories, imagined trials seems more influenced by modality-specific cognitive processes. These results provide behavioural support for the Motor-Cognitive model and contribute to understanding how movement repetition influences overt and imagined actions.

**Public Significance Statement:** Mentally rehearsing a movement is widely used in sport and in rehabilitation, but it remains unclear how closely imagined movements match real ones. We asked participants to draw, or to imagine drawing, paths around obstacles, and found that repeating a path speeded up the preparation of both real and imagined movements, whereas performance itself followed different rules in each case. Mental practice therefore supports movement planning but does not simply substitute for physical practice.

## 1. Introduction

### 1.1 Theoretical background

Motor imagery, defined as the mental simulation of a movement without physical execution (Jeannerod, 1994), is increasingly recognised as a powerful cognitive tool. Indeed, many studies have already demonstrated the benefits of motor imagery for improving performance (Toth et al., 2020) on basic and sport-related tasks (for sport see Lindsay et al., 2019; Mizuguchi et al., 2012; for basic tasks see Ruffino et al., 2017), as well as during rehabilitation for several disorders (Ietswaart et al., 2011; Jackson et al., 2001; Malouin & Richards, 2010).

It has long been assumed that motor imagery and the execution of actions rely on similar neural principles (Jeannerod, 2001). Indeed, Motor Simulation Theory proposes that imagined and executed actions should be temporally equivalent, as both share similar neural networks (Decety et al., 1989). However, the Motor Simulation Theory have been challenged by recent studies. The Motor-Cognitive model, established by Glover & Baran (2017), argues that motor imagery is more a cognitive than a motor process. This perspective is supported by behavioural observations of temporal discrepancies between imagined and executed movements, particularly when cognitive load interferes with the task (Glover & Baran, 2017) or when the actions to be performed are unfamiliar (Guillot & Collet, 2005). Consistent with these observations, a meta-analytic study conducted by Van Caenegem et al. (2026) demonstrated that the networks involved in motor imagery share greater overlap with brain areas involved in working memory than with movement execution. These temporal dissociations and neural differences raise important questions about the mechanisms underlying motor imagery and its contribution to performance improvement.

Imagery is often used as a tool for mental rehearsal; actions can be repeated in the mind, or an action can imagine prior to physical ‘repetition’ of the same movement. This repetition may induce short-term facilitation processes that influence subsequent actions in a similar way to physically repeating the same action (Smith, 1968) or repeating an action that was just observed (Edwards et al., 2003). These studies suggest that recently activated motor representations remain temporarily accessible and can influence subsequent action planning and execution. Repeating movements may therefore promote the reuse of previously successful motor strategies, a phenomenon referred to as “motor hysteresis”, which may reduce the cognitive costs associated with movement planning (Schütz & Schack, 2015). In the long-term, repeated practice typically leads to improvements in both speed and accuracy (Felix et al., 2012). Other researchers have also shown that physical practice leads to reductions in the duration of both executed and imagined movements (McAteer et al., 2025), with the amount of practice significantly impacting these effects and that combining motor imagery with physical practice produces synergistic effects on motor learning (Frank et al., 2014).

A key aspect of motor control is how the central nervous system plans movement trajectories, especially when navigating around obstacles. Van der Wel et al. (2007) demonstrated that planning curved movements induces a reaction time cost, even when obstacles are absent. This cost reflects an explicit representation of the trajectory, which persists across trials as a form of motor priming, where prior movement paths influence subsequent ones. Wong et al. (2016) further investigated this process, showing that trajectory planning operates as a distinct computational stage. Their work revealed that additional reaction time is specifically allocated to this stage when the movement path is task-relevant (e.g., avoiding barriers), independent of the movement’s curvature, and is absent in simple point-to-point reaches. These findings indicate that the brain actively represents movement paths when necessary; a process that likely engages working memory and contributes to motor imagery, where trajectories must be mentally simulated without execution. However, the question of how repetition and trajectory planning interact remains largely unexplored.

### 1.2 Aim of the study

Understanding the relationship between motor imagery and physical movement is theoretically important and has practical implications for motor performance. The present study investigated how movement repetition affects the performance of both physical and imagined movements in a barrier-constrained point-to-point reaching task, examining whether repeating a movement pattern facilitates performance across and within these two modalities. By exploring possible interactions between repetition and motor imagery in reaction times and movement times, this study aims to clarify the role of motor imagery in motor performance and its application in other various domains.

If the Motor Simulation Theory is correct, motor imagery and physical movement should rely on shared neural circuits. As a result, repetition of a movement trajectory should facilitate performance across modalities for both movement planning and execution, leading to “cross-modal” priming effects regardless of whether the movement is imagined or physically performed. In contrast, the Motor-Cognitive model posits that while imagery and physical performance may share common processes during movement preparation, they diverge during the execution phase. Consequently, repetition effects should be similar during the preparation of movements (i.e. when measuring reaction times) but may diverge during execution (i.e. when measuring imagined/real movement durations).

## 2. Methods

### 2.1 Participants

Thirty-two healthy adult participants were recruited to take part in this study. Participants were recruited from UCLouvain student community in Belgium. The sample consisted of 20 females and 12 males, 28 of them were right-handed and 4 were left-handed. The average age was 22.7 ± 3.4 years. All participants were naive to the purposes of this study and provided written informed consent before participation. Participants were compensated with 8€ for their time to complete the study (full completion was approximately 45 minutes). All participants completed the study and no participant was excluded. This experiment received ethical approval from the UCLouvain Psychological Sciences Research Ethics Committee (reference: Projet2025-94-BIS, Date: 28/11/2025).

### 2.2 Experimental design

#### 2.2.1 Set-up

Participants were seated comfortably on a chair facing the experimental set-up. The set-up comprised three vertically arranged levels (Fig. 1A). This set-up allows us to remove visual feedback of the hand during the task and allows the participant to see the path during the physical movement. The upper level consisted of a 22-inch screen (Dell, P2213t; refresh rate of 60 Hz), facing downwards, displaying the task. The middle level contained a mirror that allowed participants to view the reflected screen. The lower level housed a Wacom Intuos Pro Large graphics tablet (PTH-860-S, sampling rate 200 Hz). The tablet measured 430.4 × 287 × 8 mm, with an active drawing area of 263 × 148 mm. Task execution was performed using a wireless Wacom Pro Pen 2 stylus. The tablet recognized the stylus when it was less than 1cm from the tablet. For the task projection, since the participant was viewing the screen through a mirror, OBS Studio (version 32.0.1) was used to invert and flip the on-screen projection. This adjustment ensured that the participant perceived the display correctly, eliminating the mirror effect and allowing the task to be performed normally.

**Figure 1:**
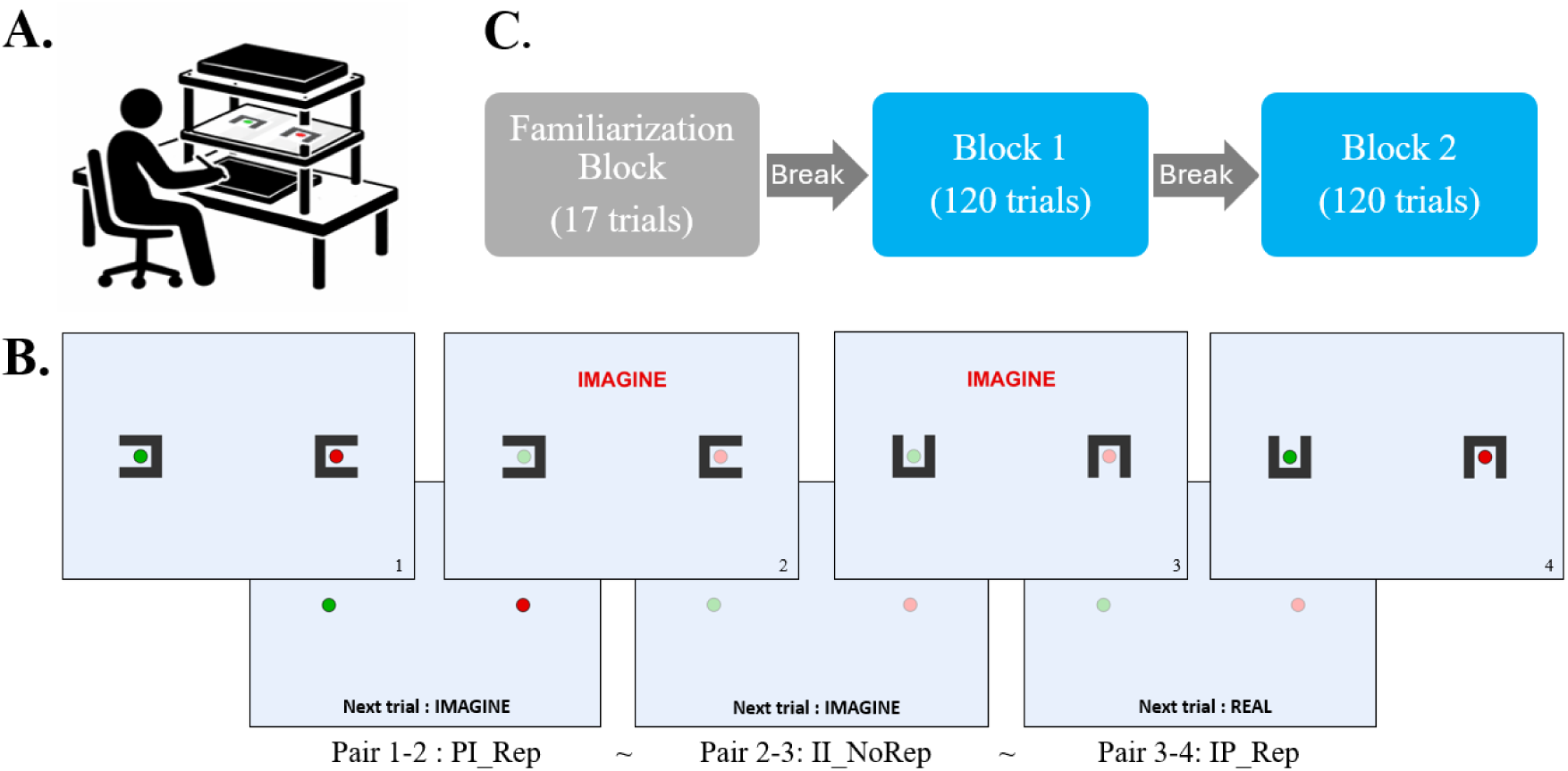
**A**. Experimental set-up **B**. Experimental design and pair creation. **C**. Overall structure

The main experimental task was coded using MATLAB (version R2024a).

#### 2.2.2 General procedure

##### 2.2.2.1 MIQ-3

Before starting the task, participants completed an electronic implementation of the French version of the Movement Imagery Questionnaire (MIQ-3) to assess their ability to perform motor imagery (for the online version see Moreno-Verdú et al., 2026; for French validation see Robin et al., 2021; for English version see Williams et al., 2012).

##### 2.2.2.2 Barrier task

The barrier task is based on a previous task developed by Wong et al. (2016).

###### Trial layout (Fig. 1B)

In each trial, a starting point (green) and a target point (red) were displayed at fixed positions on the horizontal axis of the workspace. The two points were 14 cm apart and each had a diameter of 8 mm. Barriers were displayed around the starting and the target locations, with one side open to allow the path to pass through. The barriers were 0.75 cm thick, and the opening was 1.5 cm wide. During physical trials, the green and red dots were brightly coloured. During imagined trials, the dots were still green and red, but their colours were muted. Also, to differentiate physical and imagined trials, the word ‘IMAGINE’ was displayed on the top of the screen during imagined trials with large, bold red letters.

###### Trial procedure (Fig. 1B)

Participants performed movements to connect the starting and target dots using a graphics tablet and stylus. Prior to each trial, text at the bottom of the screen informed participants as to whether the next trial would require them to physically perform or imagine performing a movement. At the start of each trial, the starting point and the target were displayed. Once the participant moved the stylus (without touching the tablet) to position the cursor inside the starting point, barriers appeared on the screen, indicating the start of the trial. In physical trials, once the barriers appeared, the participant had to press the stylus on the starting point and, as quickly and accurately as possible, draw a path to reach the target, releasing the stylus from the tablet upon reaching the end position.

In imagined trials, when the barriers appeared, participants had to click with the stylus on the starting point to indicate the start of their imagined movement, imagine the movement using a combination of kinesthetic and first-person visual imagery, and then, once they imagined reaching the target, release the stylus from the tablet. Participants were asked to maintain the same drawing speed between the physical and imagined trials (i.e. as fast and as accurately as possible). While in several cases the barriers afforded participants to take multiple different paths to reach the end position, participants were instructed to use the same real/imagined trajectory each time they saw a barrier orientation they had already experienced.

###### Overall Structure (Fig. 1C)

The experiment consisted of a familiarisation block followed by two main experimental blocks. Each block was separated by a two-minute break to avoid mental and physical fatigue.

###### Familiarization block

The familiarisation block was designed to familiarise participants with the task requirements and barrier configurations. It consisted of two trials without barriers, followed by 15 trials representing each of the 15 possible orientations that could be presented during the task (the barrier orientation where the openings faced each other was removed as it required minimal trajectory planning). During this familiarization block, all trials required the physical execution of movements.

###### Main experimental blocks

Each main block consisted of 120 trials (15 barrier orientations x 2 modalities (Physical vs Imagined) x 2 conditions (Repeated vs Non_Repeated orientation). Each orientation × modality × orientation-condition combination was presented twice, resulting in a total of 120 trials. For each barrier orientation, the trials were organised in pairs that could involve either physical movements or imagined movements, resulting in four possible combinations: physical-physical (PP), physical-imagined (PI), imagined-physical (IP) and imagined-imagined (II). In addition, the orientation of the barriers from one trial to the next could either be identical (Rep) or different (No_Rep). Trial order was pseudorandomized, with the constraint that no more than two identical barrier configurations could occur consecutively, and no rules for modalities. The second block was constructed in the same way as the first block with a different randomized order of trials following the same design. A 2-minute break was taken between the two blocks to avoid physical and/or mental fatigue.

###### Data acquisition

The position of the stylus and the movement trajectories were recorded continuously throughout each trial. To account for the differences in physical dimensions between the tablet and the screen, a calibration was performed so that one centimetre of drawing on the tablet corresponded to one centimetre on the screen. Participant reaction times and movement times were recorded for analysis. Reaction time was defined as the interval between the appearance of the barriers and the contact between the stylus and the tablet. Movement time was defined as the time elapsed between the contact between stylus and tablet on the starting point and the release of the stylus from the tablet at the end of the physical/imagined movement.

### 2.3 Data sorting and exclusion criteria

First, imagined trials involving movements greater than 4 mm (approximately 14 pixels, or the radius of the starting point) were excluded (131 trials = 1.7%). Next, all physical movements in which the participant touched either one of the two barriers were also excluded (268 trials = 3.48%). After that, certain technical issues related to the tablet or the stylus were noted. It was possible that the stylus was not recognized by the tablet when the movement was initiated, so all trials lasting less than 0.8 seconds were removed (327 trials = 4.25%). It also happened that the release of the stylus was not recognized, which resulted in the removal of all trials lasting more than 5 seconds (141 trials = 1.8%). Finally, some participants forgot to initiate the trial once the barriers appeared. All trials with a reaction time exceeding 2 seconds were also excluded from the analysis (590 trials = 7.68 %).

Finally, since our study examined the effect of previous trials on the current trial, all trials that did not have a valid previous trial were excluded from the study (1048 trials = 13.6%).

A total of 5,884 trials (72.8%) were analysed out of a maximum of 7,680 possible trials, for an average of 184 trials per participant (SD = ± 26.8), out of 240 completed trials.

### 2.4 Data analysis

No a priori power analysis was conducted. The sample size of thirty-two participants was determined on the basis of sample sizes used in previous studies employing comparable within-participant motor imagery and repetition paradigms, and of the number of observations available per participant (240 trials per participant, 120 per experimental block), which yields a large number of observations per cell for the linear mixed models. A sensitivity analysis of the effects actually detected is reported in the Strengths and Limitations section.

Statistical analyses were performed with RStudio version 2026.04.0+526 and with JASP Version 0.19.

Reaction and movement times were analysed using separate linear mixed models with a 2x2x2 design, which evaluated the effects of current trial type (physical or imagined), previous trial type (physical or imagined) and orientation (repeated or non-repeated) as well as their potential interactions. The linear mixed model was performed using the packages lmerTest in R, using the formula: *Measure (Reaction Time or Movement Time)* ∼ *Current Trial Type* × *Previous Trial Type* × *Orientation* + (1 + *Current Trial Type* + *Previous Trial Type* + *Orientation* | *Participant*). A full model with all factors and interactions in the random effect did not converge and had singularity issues, so we decided to keep the main effects without their interactions. Post hoc analyses (differences between conditions) were conducted using Bonferroni correction for multiple comparisons. Results are reported as estimated marginal means (EMM) ± standard error (SE) and alpha = 0.05 was used for statistical significance.

Pearson correlations were used to assess whether there was a link between MIQ-3 scores and the difference between physical and imagined times. Due to the lack of statistical power resulting from the small number of participants for this type of statistical analysis, the results will be provided in the supplementary materials.

## 3. Results

### 3.1 Linear Mixed Models

Full results of the 2x2x2 Linear Mixed models for reaction time and movement time are presented in Tables 1 and 2 below.

**Table 1:**
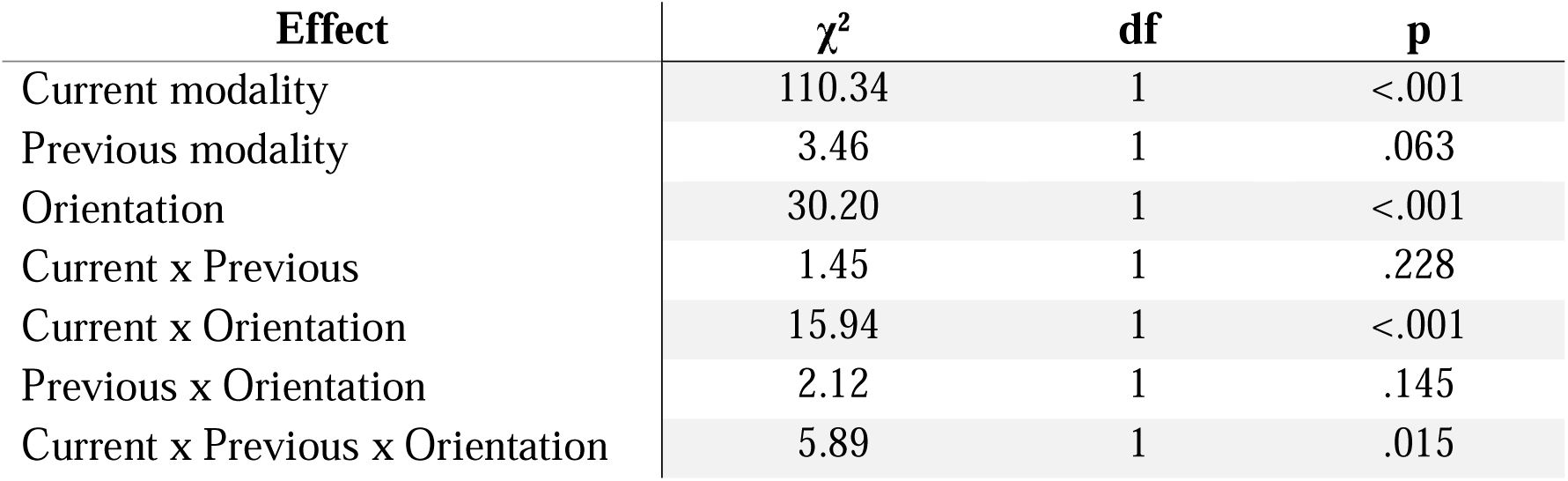
Results of the linear mixed model for reaction time.

| Effect | $\chi^2$ | df | p |
| --- | --- | --- | --- |
| Current modality | 110.34 | 1 | <.001 |
| Previous modality | 3.46 | 1 | .063 |
| Orientation | 30.20 | 1 | <.001 |
| Current x Previous | 1.45 | 1 | .228 |
| Current x Orientation | 15.94 | 1 | <.001 |
| Previous x Orientation | 2.12 | 1 | .145 |
| Current x Previous x Orientation | 5.89 | 1 | .015 |

**Table 2:** Results of the linear mixed model for movement time.

| Effect | $\chi^2$ | df | p |
| --- | --- | --- | --- |
| Current modality | 2.70 | 1 | .101 |
| Previous modality | 4.78 | 1 | .029 |
| Orientation | 1.8 | 1 | .178 |
| Current x Previous | 4.79 | 1 | .029 |
| Current x Orientation | 0.00 | 1 | .977 |
| Previous x Orientation | 1.09 | 1 | .297 |
| Current x Previous x Orientation | 0.88 | 1 | .348 |

### 3.2 Differences between current physical and imagined trials

First, regarding the Reaction Time, the analysis indicated a significant effect of the current trial type, with an overall difference between physical (*EMM = 761ms, SE = 43.6ms*) and imagined (*EMM = 1000ms, SE = 25.6ms*) conditions, whereby reaction times for physically performed trials were significantly faster than for imagined trials (*estimate = 239ms, SE = 21.6ms, z = 11.08, p < .001*) (Fig. 2A).

**Figure 2:**
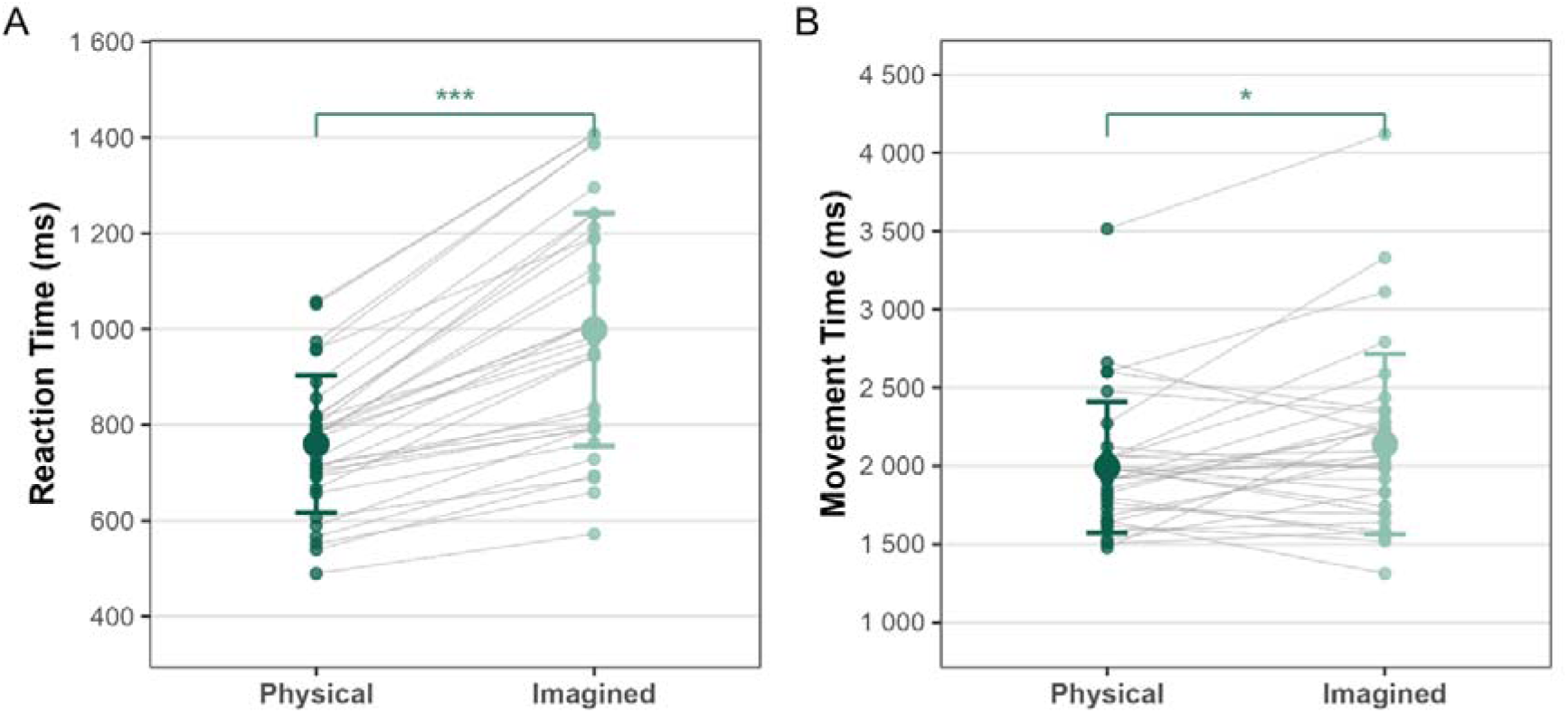
**A.** Difference between physical and imagined reaction times for current trial. **B.** Difference between physical and imagined movement times for current trial.

For movement time, the analysis also showed a significant effect of the current trial type, with an overall difference between physical (*EMM = 1991ms, SE = 74.2ms*) and imagined (*EMM = 2138ms, SE = 102ms*) movement times, such that trial durations were significantly shorter for physically performed trials compared to imagined ones (*estimate = 147ms, SE = 66.4ms, z = 2.22, p = 0.026*) (Fig. 2B).

### 3.3 Reaction Time

#### 3.3.1 Physical current trial

Post hoc analyses were performed on trials in which participants physically performed movements, in order to observe whether there was a main effect of the previous trial type (physical or imagined), orientation (repeated or non-repeated) or interactions between these factors. There was a significant effect of orientation, with shorter reaction times if the orientation was repeated compared to non-repeated (*estimate = 28.6 ms, SE = 11.3 ms, z = 2.52, p = 0.012*) (Fig. 3A). There was also a significant effect of the previous trial type, whereby physically performed trials had significantly faster reaction times if they were preceded by a physical compared to an imagined movement (*estimate = 73.9 ms, SE = 20.3 ms, z = 3.64, p = 0.0003*) (Fig. 3C). For the interaction between previous type and orientation for a current physical trial, post hoc tests indicated shorter reaction times for repeated orientations if the previous trial was physical (*estimate = 43.8ms, SE = 13.7ms, z = 3.2, p = 0.001*) but not if it was imagined (*estimate = 13.4ms, SE = 13.7ms, z = 0.98, p = 0.328*) (Fig. 3E).

**Figure 3:**
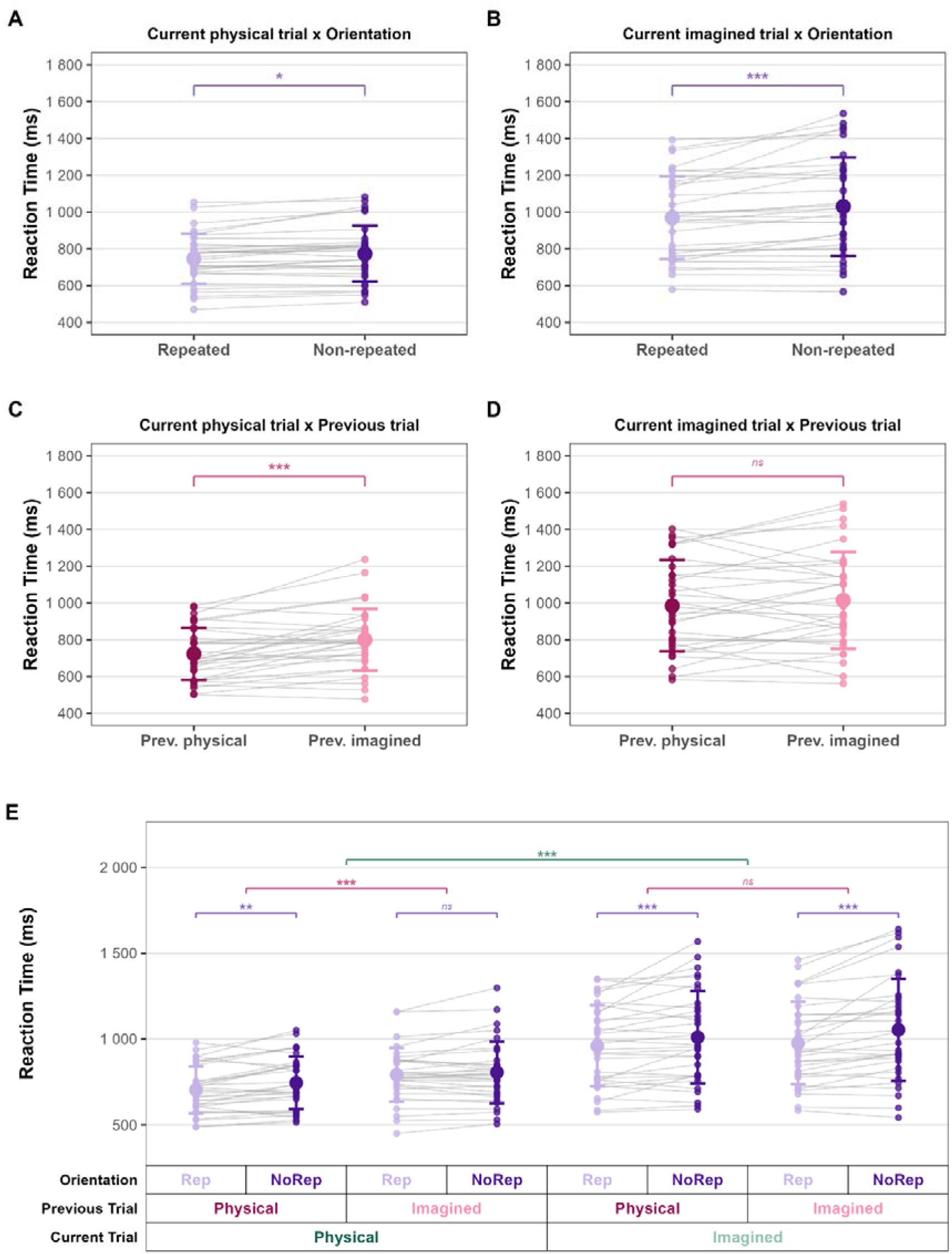
**A.** Effect of orientation for current physical trial for reaction time. **B.** Effect of orientation for current imagined trial for reaction time. **C.** Effect of previous trial type for current physical trial for reaction time. **D.** Effect of previous trial type for current imagined trial for reaction time. **E.** Interactions between current trial, previous trial type and orientation for reaction time.

#### 3.3.2 Imagined current trial

The post hoc analyses for imagined trials showed a significant main effect of orientation, whereby reaction times were shorter for repeated vs non-repeated orientations (*estimate = 63.1ms, SE = 11.2ms, z = 5.61, p < .001*) (Fig. 3B). By comparison, no significant main effect of previous trial type was present, with no statistical difference if the previous trial was physically performed or imagined (*estimate = 29.1ms, SE = 20.2ms, z = 1.44, p = 0.15*) (Fig. 3D). The post hoc tests also revealed a significant interaction effect, whereby reaction times were shorter for both previous physical (*estimate = 52ms, SE = 13.6ms, z = 3.81, p = 0.0001*) and imagined trials (*estimate = 74.1ms, SE = 13.5ms, z = 5.49, p < .001*) when the barrier orientation was repeated (Fig. 3E).

### 3.4 Movement Time

#### 3.4.1 Physical current trial

Post hoc analyses were performed to observe whether physically performed movement times were affected by the previous trial type (physical or imagined), orientation (repeated or non-repeated) or whether there was an interaction between these factors. There was a significant main effect of orientation, whereby movement times were shorter for repeated compared to non-repeated barrier orientations (*estimate = 26.6ms, SE = 12.9ms, z = 2.06, p = 0.0393*) (Fig. 4A). However, no significant effect of the previous trial type was observed (*estimate = 11ms, SE = 18ms, z = 0.61, p = 0.542*) (Fig. 4C). For the interaction between previous type and orientation for a current physical trial, post hoc tests indicated no significant interaction between conditions: the difference between repeated and non-repeated orientations was not significant for previous physical trials (*estimate = 30.3ms, SE = 18ms, z = 1.27, p = 0.203*), as well as for previous imagined trials (*estimate = 22.9ms, SE = 18ms, z = 1.68, p = 0.093*) (Fig. 4E).

**Figure 4:**
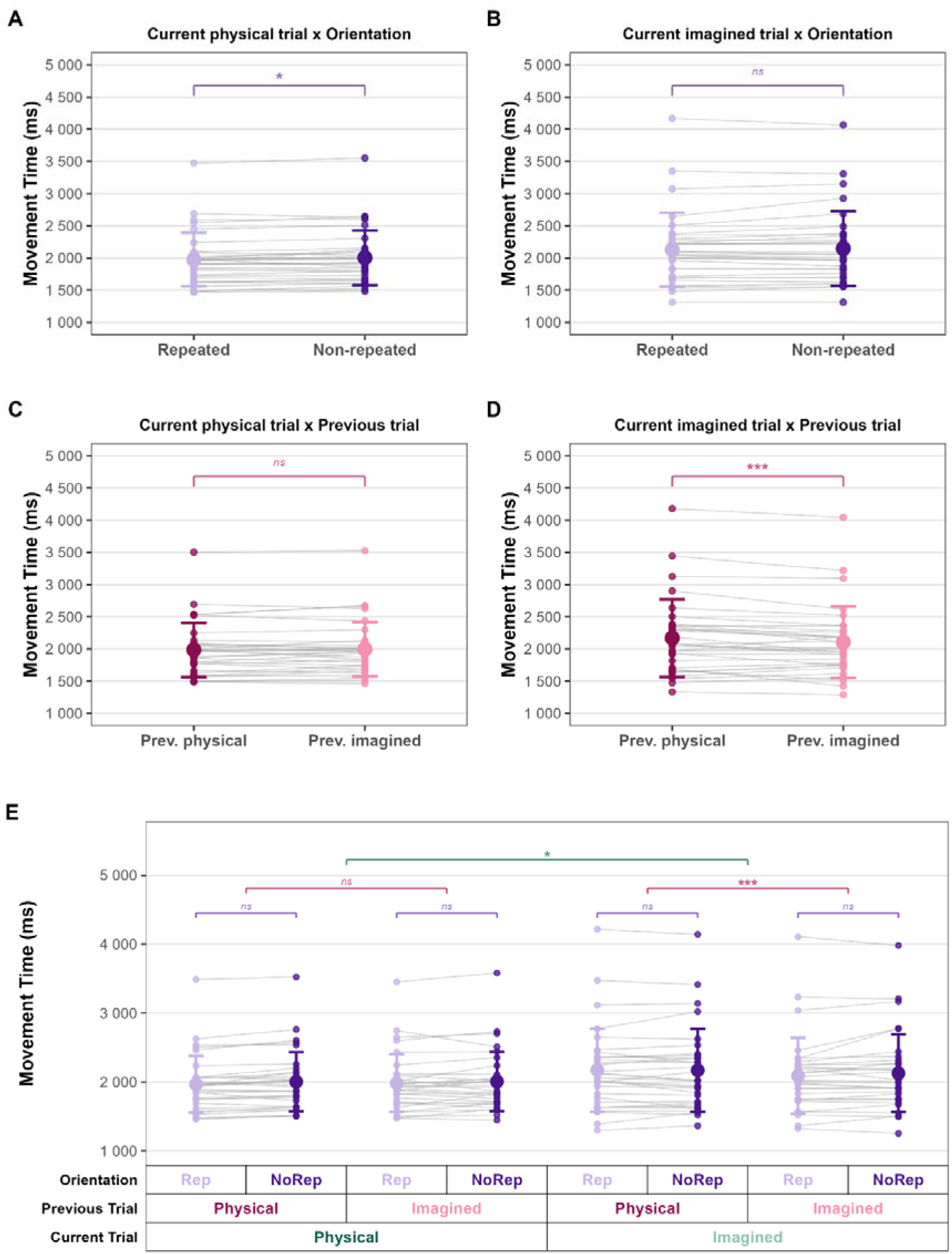
**A.** Effect of orientation for current physical trial for movement time. **B.** Effect of orientation for current imagined trial for movement time. **C.** Effect of previous trial type for current physical trial for movement time. **D.** Effect of previous trial type for current imagined trial for movement time. **E.** Interactions between current trial, previous trial type and orientation for movement time.

#### 3.4.2 Imagined current trial

For a current imagined trial, post hoc analyses didn’t reveal an effect of orientation (*estimate = 10.8ms, SE = 12.7ms, z = 0.85, p = 0.398*) (Fig. 4B). But, a main effect of previous trial type was identified, with shorter movement times if preceded by an imagined compared to a physically performed trial (*estimate = -60.4ms, SE = 17.8ms, z = -3.38, p = 0.0007*) (Fig. 4D). The interaction did not reveal a difference between repeated and non-repeated orientations in either previous physical (*estimate = -2.17ms, SE = 17.9ms, z = -0.12, p = 0.904*) nor imagined modalities (*estimate = 23.6ms, SE = 17.6ms, z = 1.35, p = 0.178*) (Fig. 4E).

## 4. Discussion

### 4.1 General overview

The goal of this study was to examine the effects of repetition (repetition of modality or repetition of orientation) on the planning and execution of real and imagined movements. An initial analysis identified that imagined movements consistently took longer than actual movements, both in terms of planning and execution (Fig. 2). Further analyses were therefore separated by whether the current movement was physically performed or imagined.

For movement preparation, indexed via reaction times, we found that repeating the same trajectory (i.e. orientation) within the same modality of movement (i.e. “physical-physical” or “imagined-imagined” trials) reduced the time required to prepare the movement. However, the presence of this effect differed when performing consecutive trials with differing modalities. Reaction times decreased when participants imagined performing a trial with the same orientation they had just physically performed (i.e. “physical-imagined” trials), but there was no corresponding benefit of imaging performing a trial prior to physically executing a trial with the same orientation (i.e. “imagined-physical” trials).

For the performance of the task, measured by movement time, physical and imagined trials were impacted in a different pattern. For physical trials, when data was pooled across all combinations of modality and orientation, trials were faster when the same orientation was repeated, through there were no modality-specific effects of the previous type of trial (i.e. no significant three-way interaction). However, imagined trials were more clearly affected by the previous trial type; imagined movements were faster if preceded by imagined trial compared to when they were preceded by a physically performed trial.

### 4.2 Physical vs Imagined trials

The longer durations of imagined compared to physical movements in the current study, for both reaction and movement times, can be considered in relation to the combined influence of task-related cognitive demands, task-switching, and task familiarity.

In relation to cognitive demands, our current findings align with recent evidence challenging the classical Motor Simulation Theory from Jeannerod (2001), which assumes a functional equivalence between motor imagery and execution. Instead, the Motor-Cognitive model proposes that motor imagery relies more on executive and working memory resources than overt action (Glover et al., 2020). Recent meta-analytic work supports this proposal, indicating that the brain network involved in motor imagery has around twice as much overlap with the brain network involved in working memory compared to that of movement execution (Van Caenegem et al., 2026). Increased cognitive load during motor imagery provides a plausible explanation for the longer movement preparation and execution durations observed in imagined movements in this study.

Another complementary explanation involves task-switching processes. In the imagery condition, participants had to alternate between a simple motor action with the stylus and the mental simulation of the movement, whereas the physical condition consisted of a continuous overt action. According to the Motor-Cognitive model, such switching places additional demands on executive control processes, which may have contributed to the longer durations observed during motor imagery.

In parallel, task familiarity and complexity could further modulate the temporal relationships between imagined and executed movements. Temporal equivalence between imagined and executed movements is typically observed for well-learned and automatized actions such as walking (Decety et al., 1989). However, in case of complex attention demanding movements or difficult tasks, the imagined movement time is often overestimated (Guillot & Collet, 2005).

In the present study, the required movement trajectory was manipulated from trial to trial with varying barrier orientations that reduced familiarity and increased planning demands. This likely led to a greater engagement of executive resources during motor imagery, which may explain the temporal discrepancies observed between imagined and executed movements.

### 4.3 Movement planning (Reaction Time)

Regarding reaction times, both physical and imagined trials revealed significant effects related to changing or maintaining the movement trajectory, suggesting that physical and imagined movements rely on partially overlapping planning mechanisms, particularly regarding the representation and reuse of movement trajectories.

Previous work has demonstrated that movement trajectories are explicitly represented during action preparation. Wong et al. (2016) showed that planning a movement path constitutes a distinct computational stage preceding movement initiation, with additional reaction time reflecting trajectory preparation rather than movement execution itself. Also, van der Wel et al. (2007) demonstrated that previously planned trajectories can influence subsequent actions, suggesting that trajectory representations remain temporarily accessible across trials. In the present study, repeating the same barrier orientation reduced the need to reconstruct a new movement path, allowing participants to reuse an already activated trajectory representation and thereby shortening reaction times.

These findings are consistent with the Motor-Cognitive model, which proposes that motor imagery and movement execution share common processes during action preparation but diverge when it is time to perform the movement (Glover & Baran, 2017). Although motor imagery is generally considered to rely more heavily on executive resources than overt action (Glover et al., 2020), both modalities appeared similarly sensitive to trajectory repetition. This suggests that imagined and executed actions rely on comparable spatial representations when preparing an upcoming movement, providing evidence for a partial functional equivalence in specific aspects of movement preparation, particularly trajectory-related representations, rather than a complete overlap of all planning processes.

However, the influence of the previous trial type differed according to the modality of the current trial. For physical movements, reaction times were shorter when the preceding trial was also physical, whereas no comparable facilitation was observed when the physical movement was preceded by an imagined movement. This suggests that, although both modalities may rely on similar trajectory representations, the information generated by a previous trial is not necessarily equally effective for preparing a subsequent overt movement.

One possible explanation is that overt execution leaves behind a sensorimotor state that is directly relevant for preparing another physical movement. This state may include recently used motor commands, sensory predictions, and proprioceptive feedback, which can facilitate the preparation of a subsequent executed action. Previous studies have shown that recently executed trajectories can bias future movement selection (van der Wel et al., 2007), while “motor hysteresis” research suggests that the motor system tends to reuse previously successful motor solutions rather than compute entirely new ones (Schütz & Schack, 2015). In contrast, motor imagery may activate or maintain a representation of the movement trajectory without generating the same sensorimotor consequences as overt execution. Thus, although an imagined movement may provide information about the trajectory itself, it may not establish the same modality-specific sensorimotor state that can be directly reused when the subsequent trial requires physical execution. Consistent with this, Roberts et al. (2024) found that the elevated trajectory required to avoid an obstacle primed the path of a subsequent executed movement when the preceding trial was executed, but not when it was imagined.

This interpretation may explain why an imagined movement immediately preceding a physical movement did not produce the same facilitation as a preceding physical movement. Importantly, this does not imply that motor imagery cannot improve subsequent physical performance in general. Rather, the present findings suggest that a single, immediately preceding imagined movement may not provide the same short-term facilitation for overt movement preparation as physical execution does. Similar effects were identified by Ramsey et al., (2010), who found that congruent imagery provided no benefit over a no-imagery baseline, an absence of facilitation they attributed to ceiling effects in simple, well-learned actions.

### 4.4 Task performance (Movement Time)

In contrast to the partial similarities observed for reaction time, movement times revealed different patterns across the modalities, suggesting distinct underlying mechanisms during task performance that depended upon whether the trial was physical or imagined.

Physical movements were influenced by orientation, with shorter durations when the same trajectory was repeated. This finding is consistent with trajectory priming accounts suggesting that recently used movement solutions remain temporarily available and can facilitate subsequent execution (van der Wel et al., 2007). Repeating the same orientation may therefore have reduced online control demands by allowing participants to reuse an already established trajectory (Wong et al., 2016).

For imagined movements, however, movement time was unaffected by orientation and was instead influenced by the modality of the previous trial. Imagined movements were completed faster when they were preceded by another imagined trial. One possible explanation is that consecutive imagery trials benefit from maintaining an imagery-specific cognitive state, whereas switching from execution to imagery requires additional cognitive reconfiguration (Glover et al., 2020). This also can be explained by the idea that transitions between execution and imagery involve inhibitory control mechanisms (Rieger et al., 2017). Furthermore, Roberts et al. (2024) found no trajectory priming following imagined trials, although in their paradigm the primed trial was always physically executed, so their findings do not directly address whether imagined movements themselves are sensitive to trajectory repetition.

### 4.5 Strengths and Limitations

This study has several strengths. First, the experimental design allowed us to distinguish between two key phases of movement, planning and execution, through the measurement of reaction and movement times. Moreover, examining these processes in both physical and imagined movements provides a perspective that has received limited attention in the literature. Finally, the design enabled the investigation of different forms of repetition, including both modality and orientation repetition.

In term of limitations, no kinematic analyses were conducted, as the focus was on behavioural measures of movement planning and movement execution rather than detailed movement characteristics. Also, only the impact of the previous trial was analysed; the present study did not examine the influence of n-2 trials or beyond, although these factors have been investigated in previous research on action simulation (Griffiths & Tipper, 2009). Although no prior power analysis was conducted, the study included 32 participants and detected several medium-to-large effects (Cohen’s d from 0.45 to 1.96 for the main significant comparisons). This suggests that the sample size was adequate for detecting the main effects of interest. However, the study may have been underpowered to detect smaller effects for some other analyses (in particular, the absence of a significant facilitation following an imagined trial should therefore be interpreted cautiously).

## 5. Conclusion

This study highlights both similarities and differences between physical and imagined movements in terms of planning and performance. Imagined movements were consistently slower than physical movements, reflecting the greater cognitive demands associated with generating and maintaining an internal representation of action.

Reaction time results suggest a partial functional equivalence between motor imagery and physical movement. The facilitation observed when the same orientation was repeated indicates that trajectory representations contribute to movement planning regardless of whether the previous trial was executed or imagined. However, only physical trials benefited from modality repetition, suggesting that overt execution may generate more stable sensorimotor representations within the same modality that can facilitate subsequent planning.

In contrast, task performance (movement time) diverged across modalities. Physical movements benefited from repeated orientation, consistent with the reuse of recently established movement solutions. Imagined movements, however, were unaffected by orientation and were instead influenced by the previous trial type, with faster performance with two consecutive imagined trials.

Overall, these results show that motor imagery and physical execution share certain mechanisms of movement preparation, particularly those related to the representation of the trajectory. However, this similarity is not complete, since the influence of the preceding movement differs depending on whether the movement is imagined or executed, and the two modalities also show differences during movement execution. These observations are consistent with the Motor-Cognitive model, which proposes common processes between imagery and execution, while suggesting the existence of mechanisms specific to each modality.

## Statements and Declarations

### Ethics approval

This experiment received ethical approval from the UCLouvain Psychological Sciences Research Ethics Committee (reference: Projet2025-94-BIS; date: 28/11/2025).

### Consent to participate

All participants provided written informed consent before participation.

## Author contributions

Elise E Van Caenegem: conceptualization, methodology, software, validation, formal analysis, investigation, resources, data curation, writing – original draft, writing – review and editing, visualization, supervision, project administration, and funding acquisition. Marcos Moreno-Verdú: formal analysis, investigation, and writing – review and editing. Charlène Truong: investigation, and writing – review and editing. Baptiste M Waltzing: investigation, and writing – review and editing. Robert M Hardwick: conceptualization, methodology, validation, investigation, resources, data curation, writing – review and editing, supervision, project administration, and funding acquisition. All authors approved the version to be published and agree to be accountable for all aspects of the work.

## Transparency Statement

We report how we determined our sample size, all data exclusions (if any), all manipulations, and all measures in the study. No participants were excluded from the reported analyses.

## Data Availability Statement

Data, analysis and visualisation scripts, are publicly available at: https://github.com/baslaboratory/Barrier-task

## Supplementary Materials

## 1. Pearson correlation (MIQ-3)

### 1.1 Results

The total MIQ-3 mean scores for all participants was 68.15 (SD ± 7.00). The correlation between the time difference between physical/imagined movements and MIQ-3 scores (Fig. 1SA) was significant (r = -0.471, p = 0.007). However, when only visual first-person perspective and kinesthetic items were considered (i.e., the modalities used to perform the task), no significant correlation was found (r = -0.218, p = 0.111) (Fig. 1SB). The average score was 43.40 (SD ± 5.67). Separate correlations were done for each modality (r_MIQ-3_K_ = -0.16, p_MIQ-3_K_ = 0.386; r_MIQ-3_V1_ = -0.29, p_MIQ-3_V1_ = 0.113, r_MIQ-3_V3_ = -0.51, p_MIQ-3_V3_ = 0.003) (Fig. 1SC,D,E).

Robust correlations were run. The points circled in red are the outliers detected by robust correlation analysis (Fig. 1S). By removing these points, coefficients of correlation are still significant for the global MIQ-3 score and no significant for specific MIQ-3 scores (r_MIQ-3 Global_ = -0.577, p_MIQ-3 Global_ = 0.001; r_MIQ-3 V1,K_ = -0.342, p_MIQ-3 V1,K_ = 0.059, r_MIQ-3_K_ = -0.221, p_MIQ-3_K_ = 0.233; r_MIQ-3_V1_ = -0.29, p_MIQ-3_V1_ = 0.113, r_MIQ-3_V3_ = -0.508, p_MIQ-3_V3_ = 0.0049).

**Figure 1S:**
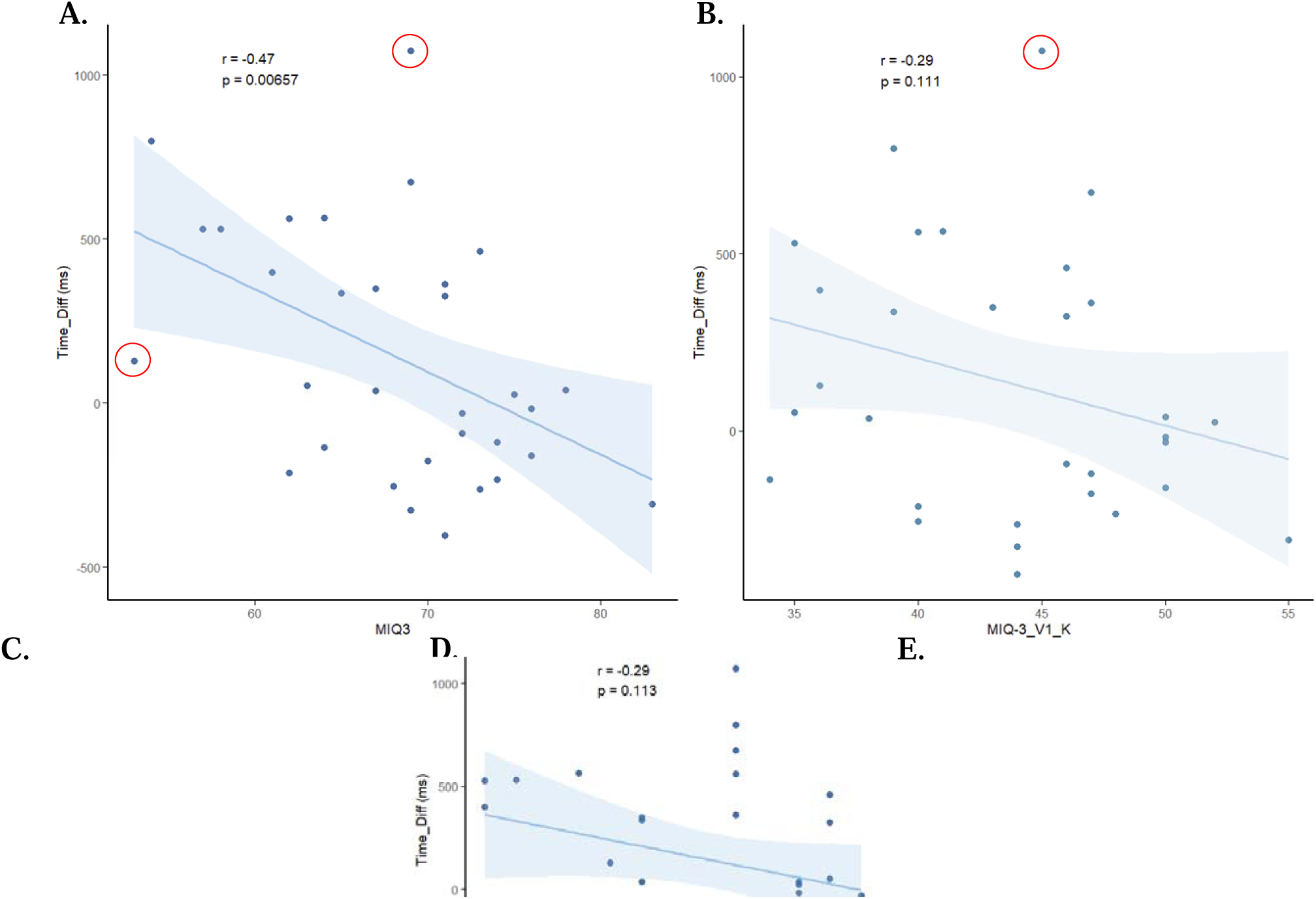
**A.** Correlation between timing differences between physical and imagined Movement Times and MIQ-3 global score. **B.** Correlation for only visual first-person perspective and kinesthetic modality. **C.** Correlation for only kinesthetic modality. **D.** Correlation for only visual first-person perspective modality. **E.** Correlation for only visual third-person perspective modality.

### 1.2 Discussion MIQ-3 correlations

Interestingly, the relationship between motor imagery ability and the timing difference between physical and imagined trials was only observed for the overall MIQ-3 score, whereas no significant association emerged for the specific imagery subscales that matched with the modality required in the task (visual first-person and kinesthetic). Motor imagery is generally considered a multimodal process involving the integration of different sensory representations (Cumming & Eaves, 2018). As a result, the specific MIQ-3 subscales may not adequately capture the combination of imagery processes involved in task performance, whereas the overall score provides a more comprehensive measure of motor imagery ability.

The absence of significant correlations for the modality-specific scores should be interpreted with caution, as correlation analyses generally require large sample sizes to reliably detect small effects. Nevertheless, the lack of association observed in the present study is consistent with previous findings. For example, Williams et al. (2015) assessed 198 participants and reported no significant relationship between MIQ-3 scores and performance on mental chronometry task.

This suggests that the absence of correlations cannot be only attributed to insufficient statistical power. Rather, it may reflect differences in the processes assessed by questionnaire-based and behavioural measures of motor imagery. The MIQ-3 was primarily designed to evaluate an individual’s capacity to generate vivid motor images (Cumming & Eaves, 2018), whereas the present task likely relied more heavily on the ability to maintain these representations over time. According to the framework proposed by Cumming & Eaves (2018), motor imagery involves several distinct cognitive processes, including image generation, maintenance, inspection, and transformation, which may not be equally captured by a single assessment tool. Supporting this interpretation, Moreno-Verdú et al. (2026) also reported no significant relationship between MIQ-3 scores and performance on the Imagined Finger Sequence Task (iFST). Together, these findings suggest that self-report questionnaires and mental chronometry tasks may assess complementary, rather than identical, components of motor imagery ability.

