## Supplementary materials for "Impact of repetition on physical and imagined movements"

1. *Pearson correlation (MIQ-3)*
   1.1 *Results*

The total MIQ-3 mean scores for all participants was 68.15 (SD ± 7.00). The correlation between the time difference between physical/imagined movements and MIQ-3 scores (Fig. 1SA) was significant (r = -0.471, p = 0.007). However, when only visual first-person perspective and kinesthetic items were considered (i.e., the modalities used to perform the task), no significant correlation was found (r = -0.218, p = 0.111) (Fig. 1SB). The average score was 43.40 (SD ± 5.67). Separate correlations were done for each modality (r_MIQ-3_K_  = -0.16, p_MIQ-3_K_ = 0.386; r_MIQ-3_V1_  = -0.29, p_MIQ-3_V1_ = 0.113, r_MIQ-3_V3_  = -0.51, p_MIQ-3_V3_ = 0.003) (Fig. 1SC,D,E).

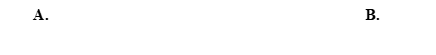
Robust correlations were run. The points circled in red are the outliers detected by robust correlation analysis (Fig. 1S). By removing these points, coefficients of correlation are still significant for the global MIQ-3 score and no significant for specific MIQ-3 scores (r_MIQ-3 Global_  = -0.577, p_MIQ-3 Global_ = 0.001; r_MIQ-3 V1,K_  = -0.342, p_MIQ-3 V1,K_ = 0.059, r_MIQ-3_K_  = -0.221, p_MIQ-3_K_ = 0.233; r_MIQ-3_V1_  = -0.29, p_MIQ-3_V1_ = 0.113, r_MIQ-3_V3_  = -0.508, p_MIQ-3_V3_ = 0.0049).

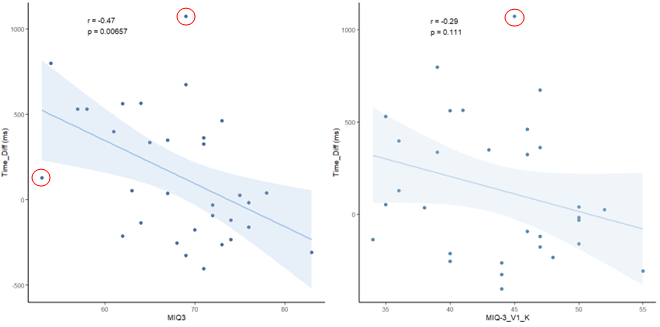

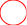

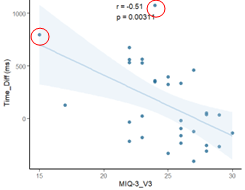

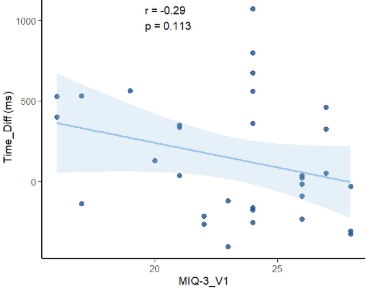

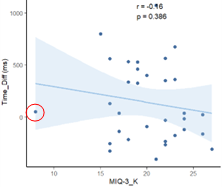

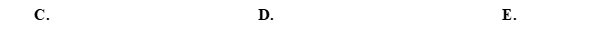

*Figure 1S:* **A.** Correlation between timing differences between physical and imagined Movement Times and MIQ-3 global score. **B.** Correlation for only visual first-person perspective and kinesthetic modality. **C.** Correlation for only kinesthetic modality. **D.** Correlation for only visual first-person perspective modality. **E.** Correlation for only visual third-person perspective modality.
